# Reproductive period, endogenous estrogen exposure and dementia incidence among women in Latin America and China; a 10/66 population-based cohort study

**DOI:** 10.1101/137687

**Authors:** Martin J Prince, Daisy Acosta, Mariella Guerra, Yueqin Huang, Ivonne Z Jimenez-Velazquez, Juan J Llibre Rodriguez, Aquiles Salas, Ana Luisa Sosa, Kia-Chong Chua, Michael E Dewey, Zhaorui Liu, Rosie Mayston, Adolfo Valhuerdi

**Author notes:** Email addresses: MARTIN J PRINCE, DAISY ACOSTA, MARIELLA GUERRA, YUEQIN HUANG, IVONNE Z JIMENEZ-VELAZQUEZ, JUAN J LLIBRE RODRIGUEZ, AQUILES SALAS, ANA LUISA SOSA, KIA-CHONG CHUA, MICHAEL E DEWEY, ZHAORUI LIU, ROSIE MAYSTON, ADOLFO VALHUERDI. Corresponding author (MJP).

## Abstract

**Background:** Exposure to endogenous estrogen may protect against dementia, but evidence remains equivocal. Such effects may be assessed more precisely in settings where exogenous estrogen administration is rare. We aimed to determine whether reproductive period (menarche to menopause), and other indicators of endogenous estrogen exposure are inversely associated with dementia incidence

**Methods:** Population-based cohort studies, of women aged 65 years and over in urban sites in Cuba, Dominican Republic Puerto Rico and Venezuela, and rural and urban sites in Peru, Mexico and China. Sociodemographic and risk factor questionnaires were administered to all participants, including ages at menarche, birth of first child, and menopause, and parity, with ascertainment of incident 10/66 dementia, and mortality, three to five years later.

**Results:** 9,428 women participated in the baseline phase, with response rates of 72–98% by site. The ‘at risk’ cohort comprised 8,466 dementia-free women. Mean age at baseline varied from 72.0 to 75.4 years, lower in rural than urban sites and in China than in Latin America. Mean parity was 4.1, generally higher in rural than urban sites and ranging from 2.4-7.2 by site. 6,854 women with baseline reproductive period data were followed up for 26,463 person years. There were 692 cases of incident dementia, and 895 dementia free deaths. Pooled meta-analysed fixed effects, per year, for reproductive period (Adjusted Sub-Hazard Ratio [ASHR] 1.001, 95% CI 0.988-1.015) did not support any association with dementia incidence, with no evidence also for any interaction between APOE genotype and reproductive period (ASHR 0.993, 95% CI 0.947-1.041, I^2^=39.3%). Similarly no association was observed between incident dementia and; ages at menarche, birth of first child, and menopause: nulliparity; or index of cumulative endogenous estrogen exposure. Greater parity was positively associated with incident dementia (ASHR 1.030, 95% CI 1.002-1.059, I^2^=0.0%).

**Conclusions:** We found no evidence to support the theory that natural variation in cumulative exposure to endogenous oestrogens across the reproductive period influences the incidence of dementia in late life.

## Introduction

Estrogen exerts potentially helpful effects on brain synapse structure and function in regions such as the prefrontal cortex and hippocampus[1]. In women, endogenous estrogen exposure (EEE) occurs mainly during the reproductive phase. Estrogen levels rise during pregnancy, but fall postnatally, particularly with breastfeeding, and are lower after a first pregnancy than in nulliparous women. Earlier menarche and later menopause (hence longer reproductive period), nulliparity or lower parity, older age at birth of first child, and less breastfeeding are therefore proxy indicators of lifetime EEE[2].

The hypothesis that estrogen is neuroprotective for women is supported by inverse associations between indicators of lifetime EEE and late-life cognitive function[2–7], and prospective and historic cohort studies indicating adverse cognitive outcomes associated with premature surgically-induced menopause[8], and premature ovarian failure (POF)[9]. However, the evidence remains inconclusive. Only two studies of EEE were population-based, effects on cognition were small, and not always replicated[10]. Although effect sizes linked to oopherectomy and POF are larger[8,9], associations with dementia were not replicated in a large Finland registry linkage study[11].

Few studies have examined the effects of EEE on cognitive decline, incident dementia or Alzheimer’s disease (AD). In the population-based Esprit study in France, EEE indicators were associated in the hypothesized direction with baseline cognitive function, but not with cognitive decline over the next four years[4]. In case-control studies, childlessness was inversely associated with AD among women but not men[12], and increasing numbers of pregnancies were associated with AD, and age of onset among cases[13]. In a nested case-control study, AD risk increased with increasing age at menarche[14]. The largest and most definitive study to date was carried out in the population-based Rotterdam cohort; 3601 postmenopausal women aged 55 years or older were followed up for a median of 6.3 years (21,046 person years)[15]. Counter to the hypothesis, women with natural menopause and more reproductive years had an increased risk of dementia (adjusted RR for highest versus lowest quarter 1.78, 95% CI confidence interval [CI] 1.12-2.84). The association was modified by APOE genotype, with a stronger association among APOE e4 carriers, while among non-carriers no association with dementia or AD was observed.

We set out to study associations between indicators of EEE and dementia incidence in the 10/66 Dementia Research Group’s population-based cohort studies in seven urban and three rural catchment area sites in six Latin American countries, and China. Historically, these populations were characterised by higher fertility rates, and a greater variation in age at first birth and parity than in high income country populations. Contemporary market penetration data suggest very low rates of use of hormone replacement therapy (exogenous estrogen)[16], allowing the effects of EEE to be estimated more precisely. Our primary hypothesis is that a longer reproductive period is independently associated with a lower risk of incident dementia. Our secondary hypotheses are that younger age at menarche, older age at menopause, lower parity, older age at birth of first child, and higher indices of cumulative endogenous estrogen exposure (ICEEE), are each associated with a lower risk of incident dementia. Finally, we test the hypothesis that APOE genotype modifies any effect of reproductive period on incident dementia risk.

## Materials and Methods

The 10/66 population-based study protocols for baseline and incidence waves [17], and a full description of the cohort profile [18] are available in open access publications. Relevant details are provided here. One-phase population-based surveys were carried out of all residents aged 65 years and over in geographically defined catchment areas (urban sites in Cuba, Dominican Republic, Puerto Rico and Venezuela, and urban and rural sites in Mexico, Peru, and China)[17]. Baseline surveys were completed between 2003 and 2007, other than in Puerto Rico (2007-2009). The target sample was 2000 for each country, and 3000 for Cuba. The baseline survey included clinical and informant interviews, and physical examination. DNA collections were carried out in the Latin American countries, and APOE genotype determined in Cuba, Dominican Republic, Venezuela and Puerto Rico. Incidence waves were subsequently completed, with a mortality screen, between 2007 and 2011 (2011-2013 in Puerto Rico) aiming for 3-4 years follow-up in each site[19]. Assessments were identical to baseline protocols for dementia ascertainment, and similar in other respects. We revisited participants’ residences on up to five occasions. When no longer resident we sought information on their vital status and current residence, from additional contacts recorded at baseline. Where participants had moved away, we sought to re-interview them, even outside the catchment area. If deceased, we recorded the date, and completed an informant verbal autopsy, including evidence of cognitive and functional decline suggestive of dementia onset between baseline assessment and death[20].

### Measures

The 10/66 population-based study interview covers dementia diagnosis, mental disorders, physical health, anthropometry, demographics, an extensive risk factor questionnaire, disability, health service utilisation, care arrangements and strain[17].

Only relevant assessments are detailed here.

Reproductive history measures: Evidence suggests that longer reproductive period (age at menopause minus age at menarche), lower parity, older age at birth of first child, and greater postmenopausal body mass are indicators of higher EEE across the life course [2,5]. Age at menarche, age at menopause, number of live births, and age at birth of first child were ascertained from all female participants at baseline interview, using four questions

1. How old were you when your periods began?
2. How many children did you have?
3. How old were you when your first child was born?
4. How old were you when you had the first symptoms of the menopause?

Weight was not measured at baseline, so waist circumference (in centimetres) was used as the relevant proxy indicator instead. Following the method proposed by Smith et al[2] each indicator was z-scored, and a composite Index of Cumulative Endogenous Estrogen Exposure (ICEEE) calculated as ((age at menopause + age at birth of first child + waist circumference)-(age at menarche + number of children).

Confounders and other covariates: Age, education, marital status, household assets, tobacco consumption (ever versus never), and hazardous drinking (>21 units per week, before the age of 65) were all ascertained in the baseline questionnaire. Height, leg length, skull circumference and waist circumference were measured in the physical examination.

Dementia: 10/66 dementia diagnosis is allocated to those scoring above a cutpoint of predicted probability for dementia, calculated using coefficients from a logistic regression equation developed, calibrated and validated cross-culturally in the 25 centre 10/66 pilot study[21], applied to outputs from a) a structured clinical interview, the Geriatric Mental State[22], b) two cognitive tests; the Community Screening Instrument for Dementia (CSI-D) COGSCORE[23] and the modified Consortium to Establish a Registry for Alzheimer’s Disease (CERAD) 10 word list learning task with delayed recall[24], and c) informant reports of cognitive and functional decline from the CSI-D RELSCORE[23].

The criterion, concurrent and predictive validity of the 10/66 diagnosis were superior to that of the DSM-IV criterion in subsequent evaluations[25–28]. For those who died between baseline and follow-up we diagnosed ‘probable incident dementia’ by applying three criteria:

1. A score of more than two points on the RELSCORE, from the post-mortem informant interview, with endorsement of either ‘deterioration in memory’ or ‘a general deterioration in mental functioning’ or both, and
2. an increase in RELSCORE of more than two points from baseline, and
3. the onset of these signs noted more than six months prior to death.

In the baseline survey, the first criterion would have detected those with either DSM-IV or 10/66 dementia with 73% sensitivity and 92% specificity[20].

The prevalence[25] and incidence[20] of 10/66 dementia in the current cohorts have been reported.

### Analyses

We used release 2.0 of the 10/66 dementia incidence data archive (October 2015), and STATA version 11 for all analyses.

For each site we

1. describe participants’ status at follow-up, age at menarche and menopause, and reproductive period for all those included in the cohort analysis (reinterviewed or deceased).
2. describe cohort characteristics by quarters of reproductive period, with tests for linear trends (one-way ANOVA or Chi-squared tests for trend, as appropriate).
3. model the effect of ages at menarche, birth of first child, and menopause; reproductive period; parity; ICEEE; and premature ovarian failure on 10/66 dementia incidence using a competing-risks regression derived from Fine and Gray’s proportional subhazards
4. model [29] (Stata stcrreg command), based on a cumulative incidence function, indicating the probability of failure (dementia onset) before a given time, acknowledging the possibility of a competing event (dementia-free death). Time to death was the time from baseline interview to the exact date of death. Time to dementia onset (which could not be ascertained precisely was the midpoint between baseline and follow-up interview. Competing risks regression keeps those who experience competing events at risk so that they can be counted as having no chance of failing. We report adjusted sub-hazard ratios (ASHR) with robust 95% confidence intervals adjusted for household clustering.

We also test for modification by APOE genotype of the effect of reproductive period on dementia incidence, by extending the adjusted models described above by the appropriate interaction terms, and also by restricting the analysis to those with no APOE e4 genotype. We fit all models separately for each site and combine them using a fixed effects meta-analysis. Higgins I^2^ estimates the proportion of between-site variability in the estimates accounted for by heterogeneity, as opposed to sampling error; up to 40% heterogeneity is conventionally considered negligible, while up to 60% reflects moderate heterogeneity [30].

The study protocol and the consent procedures were approved by the King’s College London research ethics committee and in all countries where the research was carried out: 1-Medical Ethics Committee of Peking University the Sixth Hospital (Institute of Mental Health, China); 2-the Memory Institute and Related Disorders (IMEDER) Ethics Committee (Peru); 3-Finlay Albarran Medical Faculty of Havana Medical University Ethical Committee (Cuba); 4-Hospital Universitario de Caracas Ethics Committee (Venezuela); 5-Consejo Nacional de Bioética y Salud (CONABIOS, Dominican Republic); 6-Instituto Nacional de Neurología y Neurocirugía Ethics Committee (Mexico); 7-University of Puerto Rico, Medical Sciences Campus Institutional Review Board (IRB)

## Results

### Sample characteristics

In all, 9,428 interviews were completed with women, at baseline, in the 10 sites in seven countries. Response proportions at baseline varied between 72% and 98%, and exceeded 80% in all sites other than urban China[25]. The ‘at risk’ cohort comprised 8,466 dementia-free women (Table 1). Mean age at baseline ranged from 72.0 to 75.4 years, lower in rural than urban sites and in China than in Latin America. Educational levels were lowest in rural China (84% not completing primary education), rural Mexico (83%), Dominican Republic (73%), and urban Mexico (57%) and highest in urban Peru (11%), Puerto Rico (23%), and Cuba (26%). In other sites, between one-third and one-half of participants had not completed primary education. Seven percent of women were nulliparous (from 0.4% in rural Peru to 14.6% in Cuba), strongly associated with never having been married (34% of never married women and 5% of married women were nulliparous). Mean parity was 4.1 (SD 3.0), higher in rural than urban sites and ranging from 2.4 (urban Cuba) to 7.2 (rural Mexico). There was significant between site variation in ages at menarche and menopause, and reproductive period. Menarche (R^2^=15.7%) was earlier in Latin American and urban sites. Menopause (R^2^=4.6%) was later in Chinese sites. Reproductive period (R^2^=1.6%) showed no clear pattern of variation among sites. From the ‘at risk’ cohort, 1,451 participants (17.1%) were lost to follow-up; 6,854 women with baseline reproductive period data were followed up for 26,463 person years (Table 1).

**Table 1.** Cohort flow, by site

| Site | At risk (n) | Status at follow up |  | Age at menarche and menopause |  | Reproductive period |  | Outcome (menarche cohort analysis) |  |
| --- | --- | --- | --- | --- | --- | --- | --- | --- | --- |
|  |  | Status | Number (%) | Menarche Mean (SD) | Menopause Mean (SD) | Mean (SD) | Numbers in cohort analysis (pyears) | Outcome | Number (%) |
| Cuba | 1628 | Interviewed | 1249 (76.7%) | 12.7 (1.8)<br>MV=7 | 48.0 (5.9)<br>MV=16 | 35.3 (6.1)<br>MV=18 | 1473 (6028) | Censored | 1101 (74.7%) |
|  |  | Deceased | 255 (15.7%) |  |  |  |  | Incident dementia | 127 (8.6%) |
|  |  | Lost | 124 (7.6%) |  |  |  |  | Competing risk | 245 (16.6%) |
| Dominican Republic | 1156 | Interviewed | 736 (63.7%) | 13.6 (2.0)<br>MV=37 | 46.3 (7.2)<br>MV=66 | 32.7 (7.3)<br>MV=81 | 911 (3945) | Censored | 620 (68.1%) |
|  |  | Deceased | 212 (18.3%) |  |  |  |  | Incident dementia | 113 (12.4%) |
|  |  | Lost | 208 (18.0%) |  |  |  |  | Competing risk | 178 (19.5%) |
| Puerto Rico | 1183 | Interviewed | 818 (69.1%) | 13.0 (1.9)<br>MV=18 | 45.7 (7.3)<br>MV=29 | 32.8 (7.5)<br>MV=37 | 905 (3686) | Censored | 712 (78.7%) |
|  |  | Deceased | 118 (10.0%) |  |  |  |  | Incident dementia | 97 (10.7%) |
|  |  | Lost | 247 (20.9%) |  |  |  |  | Competing risk | 96 (10.6%) |
| Peru Urban | 802 | Interviewed | 540 (67.3%) | 12.9 (1.7)<br>MV=15 | 45.7 (6.1)<br>MV=17 | 32.9 (6.0)<br>MV=25 | 548 (1544) | Censored | 496 (90.5%) |
|  |  | Deceased | 26 (3.2%) |  |  |  |  | Incident dementia | 27 (4.9%) |
|  |  | Lost | 236 (29.5%) |  |  |  |  | Competing risk | 25 (4.6%) |
| Peru Rural | 271 | Interviewed | 211 (77.9%) | 13.0 (1.8)<br>MV=7 | 46.6 (5.9)<br>MV=15 | 33.6 (6.0)<br>MV=17 | 221 (757) | Censored | 185 (83.7%) |
|  |  | Deceased | 19 (7.0%) |  |  |  |  | Incident dementia | 19 (8.6%) |
|  |  | Lost | 41 (15.1%) |  |  |  |  | Competing risk | 17 (7.7%) |
| Venezuela | 1150 | Interviewed | 769 (66.9%) | 13.1 (1.6)<br>MV=5 | 48.5 (5.6)<br>MV=50 | 35.4 (5.7)<br>MV=51 | 844 (3339) | Censored | 674 (79.9%) |
|  |  | Deceased | 80 (7.0%) |  |  |  |  | Incident dementia | 101 (12.0%) |
|  |  | Lost | 301 (26.1%) |  |  |  |  | Competing risk | 69 (8.2%) |
| Mexico Urban | 597 | Interviewed | 460 (77.1%) | 13.4 (1.6)<br>MV=10 | 47.1 (5.9)<br>MV=27 | 33.7 (6.2)<br>MV=30 | 494 (1430) | Censored | 423 (85.6%) |
|  |  | Deceased | 44 (7.4%) |  |  |  |  | Incident dementia | 32 (6.5%) |
|  |  | Lost | 93 (15.5%) |  |  |  |  | Competing risk | 39 (7.9%) |
| Mexico Rural | 547 | Interviewed | 411 (75.1%) | 14.0 (1.5)<br>MV=20 | 45.6 (5.6)<br>MV=30 | 31.6 (5.7)<br>MV=40 | 439 (1221) | Censored | 350 (79.7%) |
|  |  | Deceased | 48 (8.8%) |  |  |  |  | Incident dementia | 46 (10.5%) |
|  |  | Lost | 88 (16.1%) |  |  |  |  | Competing risk | 43 (9.8%) |
| China Urban | 609 | Interviewed | 419 (68.8%) | 14.8 (1.7)<br>MV=0 | 48.2 (3.4)<br>MV=0 | 33.5 (3.8)<br>MV=0 | 496 (2252) | Censored | 368 (74.2%) |
|  |  | Deceased | 77 (12.6%) |  |  |  |  | Incident dementia | 62 (12.5%) |
|  |  | Lost | 113 (19.6%) |  |  |  |  | Competing risk | 66 (13.3%) |
| China Rural | 523 | Interviewed | 394 (75.3%) | 15.3 (1.2)<br>MV=0 | 50.3 (2.5)<br>MV=2 | 35.0 (2.8)<br>MV=2 | 523 (2263) | Censored | 338 (64.6%) |
|  |  | Deceased | 129 (24.7%) |  |  |  |  | Incident dementia | 68 (13.0%) |
|  |  | Lost | 0 (0.0%) |  |  |  |  | Competing risk | 117 (22.4%) |
| Total | 8466 | Interviewed | 6007(71.0%) | 13.4 (1.9)<br>MV=119 | 47.3 (6.1)<br>MV=252 | 33.9 (6.2)<br>MV=301 | 6854 (26463) | Censored | 5267 (76.8%) |
|  |  | Deceased | 1008 (11.9%) |  |  |  |  | Incident dementia | 692 (10.1%) |
|  |  | Lost | 1451 (17.1%) |  |  |  |  | Competing risk | 895 (13.1%) |

In the at risk cohort, longer reproductive period was associated, predictably, with earlier menarche and (particularly) later menopause, and with a higher ICEEE (Table 2). Longer reproductive period was also associated with older age; with being more likely to complete primary education; with lower parity; with older age at birth of first child; with a lower prevalence of hazardous drinking and stroke; with taller stature, and with better baseline cognitive function (the composite CSI-D COGSCORE, and the CERAD animal naming test, but not the CERAD 10 word list delayed recall). The effect on COGSCORE (effect size per quarter of reproductive period +0.064, 95% CI +0.021 to +0.107, r^2^=0.1%) was no longer statistically significant having adjusted for age, education, marital status, stroke, hazardous alcohol use, and height (+0.024, 95% CI-0.019 to +0.067, r^2^=0.0%). The effect on animal naming (+0.195, 95% CI +0.094 to +0.295, r^2^=0.2%) also lost statistical significance after adjusting for the same covariates (+0.092, 95% CI-0.006 to +0.190, r^2^=0.0%). Although the proportion with one or more APOE e4 alleles did decline significantly across quarters of reproductive period (p=0.048, Table 2), neither mean reproductive period (p=0.08), nor mean age at menarche (p=0.38), nor mean age at menopause (p=0.17) differed significantly across APOE genotypes.

**Table 2.** Cohort characteristics at baseline, by quarters of reproductive period

| Reproductive period (range in years)<br>(MV=366) | 1<sup>st</sup> quarter<br>(<31)<br>n=2050 | 2<sup>nd</sup> quarter<br>(31-34)<br>n=1827 | 3<sup>rd</sup> quarter<br>(35-37)<br>n=1948 | 4<sup>th</sup> quarter<br>(>38)<br>n=2275 | All combined<br>n=8466 | Test for linear<br>trend |
| --- | --- | --- | --- | --- | --- | --- |
| Mean age in years (SD) | 73.5 (6.5) | 73.4 (6.4) | 73.8 (6.7) | 73.9 (6.9) | 73.8 (6.7) | 5.8, 0.02 |
| Did not complete primary education (%) | 872 (42.6%) | 825 (44.2%) | 897 (46.0%) | 803 (35.4%) | 2613 (42.8%) | 18.5, <0.001 |
| Mean age at menarche | 13.7 (2.1) | 13.7 (1.9) | 13.7 (1.5) | 12.6 (1.7) | 13.4 (1.9) | 394.2, <0.001 |
| Mean age at menopause | 39.2 (4.6) | 46.4 (2.2) | 49.6 (1.6) | 53.1 (2.9) | 47.3 (6.1) | 22652.8, <0.001 |
| Mean age at birth of first child | 22.0 (4.9) | 22.3 (4.6) | 22.4 (5.1) | 22.6 (5.4) | 22.3 (5.0) | 8.4, 0.004 |
| Nulliparous (%) | 173 (8.6%) | 103 (5.7%) | 103 (5.4%) | 159 (7.1%) | 576 (6.9%) | 3.5, 0.06 |
| Mean live births (SD) | 4.2 (3.1) | 4.2 (2.9) | 4.1 (2.9) | 3.9 (2.9) | 4.1 (3.0) | 14.7, <0.001 |
| Index of estrogen exposure (n=6504) | -1.66 (2.13) | -0.44 (1.90) | +0.24 (1.99) | +1.59 (2.16) | -0.01 (2.38) | 2194.6, <0.001 |
| Ever smoked (%) | 489 (23.9%) | 386 (21.2%) | 374 (19.3%) | 567 (25.0%) | 1907 (22.6%) | 0.3, 0.59 |
| Hazardous drinker (%) | 78 (4.0%) | 41 (2.4%) | 31 (1.7%) | 51 (2.5%) | 219 (2.8%) | 10.1, 0.002 |
| Stroke (%) | 136 (6.6%) | 104 (5.7%) | 84 (4.3%) | 122 (5.4%) | 464 (5.5%) | 5.0, 0.03 |
| Diabetes (%) | 398 (19.5%) | 337 (18.4%) | 325 (16.7%) | 443 (19.5%) | 1571 (18.6%) | 0.1, 0.76 |
| Obesity (%) | 1145 (59.3%) | 959 (54.9%) | 965 (52.2%) | 1283 (60.9%) | 4352 (57.0%) | 0.4, 0.53 |
| Mean height in centimetres | 153.0 (7.5) | 153.3 (7.2) | 154.1 (7.3) | 153.9 (7.3) | 153.5 (7.4) | 20.6, <0.001 |
| Mean leg length in centimetres | 85.1 (6.9) | 84.8 (6.9) | 84.4 (7.9) | 85.5 (7.1) | 84.9 (7.2) | 1.7, 0.18 |
| Mean skull circumference in centimetres | 54.9 (2.1) | 54.8 (2.2) | 54.9 (2.2) | 54.8 (2.3) | 54.8 (2.2) | 0.0, 0.90 |
| Mean CSI-D COGSCORE | 30.2 (2.3) | 30.3 (2.3) | 30.3 (2.4) | 30.4 (2.2) | 30.2 (2.4) | 8.4, 0.004 |
| Mean CERAD 10 word recall | 4.9 (2.1) | 5.0 (2.1) | 4.8 (2.0) | 5.0 (2.0) | 4.9 (2.0) | 0.2, 0.64 |
| Mean CERAD animal naming | 15.4 (5.1) | 15.5 (5.1) | 15.4 (5.3) | 16.0 (5.6) | 15.5 (5.3) | 14.4, <0.001 |
| One or more APOE e4 alleles<sup>1</sup> (%) | 188 (22.3%) | 111 (18.5%) | 151 (20.1%) | 224 (18.2%) | 709 (19.9%) | 3.9, 0.05 |

1. This analysis was conducted using data from the four sites where APOE genotype was available; Cuba (n=1427), Dominican Republic (n=635), Venezuela (n=599), and Puerto Rico (n=907), total (n= 3568)

### Primary hypothesis-reproductive period and incident dementia

Controlling for age, education and household assets, there was no association between reproductive period and dementia incidence, either in individual sites or after pooled metaanalysis (ASHR per year 1.001, 95% CI 0.988-1.015, I^2^ 0.0%) (Table 3). The effect of reproductive period was unchanged after controlling additionally for marital status, hazardous alcohol use, stroke, and height (ASHR per year 1.001, 95% CI 0.987-1.015, I^2^ 0.0%). In Cuba, Dominican Republic, Venezuela and Puerto Rico, where APOE genotype was available, there was no evidence for an interaction between APOE genotype and reproductive period (ASHR 0.993, 95% CI 0.947-1.041, I^2^=39.3%), or for an effect of reproductive period among non-carriers of the e4 allele (ASHR 1.012, 95% CI 0.991-1.034, I^2^=0.0%). Controlling for APOE genotype in addition to age, education, household assets, marital status, hazardous alcohol use, stroke and height did not affect the fully adjusted association between reproductive period and incident dementia (ASHR 1.018, 95% CI 0.998-1.038, I^2^=0.0%),

#### Secondary hypotheses

Controlling for age, education and household assets, there was no association between dementia incidence and either age at menarche (pooled ASHR per year 0.986, 95% CI 0. 944-1.030, I^2^ 0.0%), or age at menopause (ASHR per year 1.000, 95% CI 0.986-1.013, I^2^ 0.0%) (Table 3). There was also no association between premature ovarian failure (before the age of 40 years) and incident dementia (ASHR 1.19, 95% CI 0.91-1.55, I^2^ 0. 0%). However, as hypothesized, greater parity was associated with incident dementia, controlling for marital status as well as age, education and assets (ASHR per birth 1.030, 95% CI 1.002-1.059, I^2^ 0.0%). The effect of nulliparity, a rare exposure, was difficult to estimate with no exposed incident cases in urban and rural Peru or rural China; in the remaining sites there was no association (ASHR 1.16, 95% CI 0.86-1.56, I^2^ 30.2%). Neither was there any evidence for an association between the ICEEE and incident dementia (ASHR per SD 0.987, 95% CI 0.951-1.025, I^2^ 0.0%).

**Table 3.**
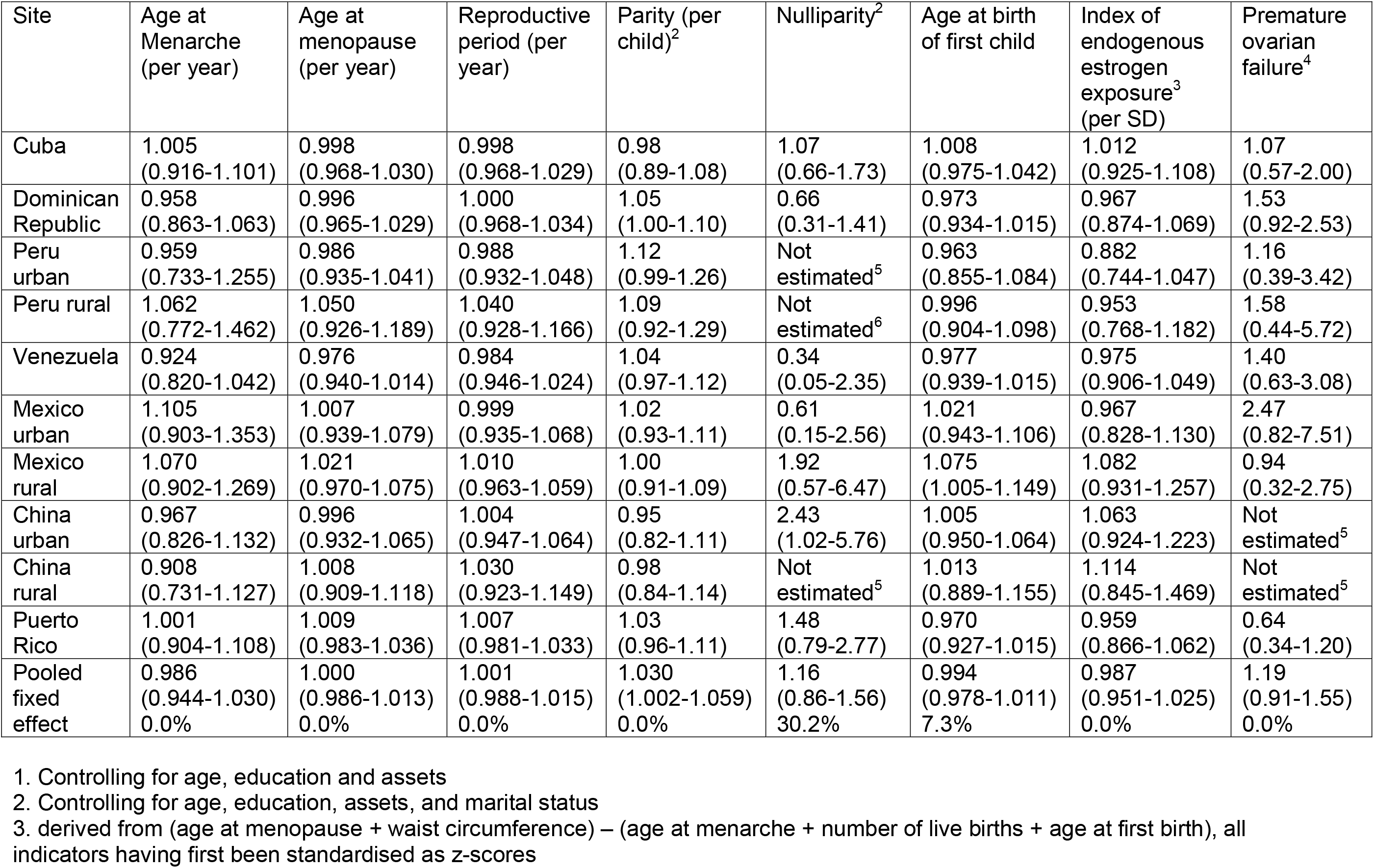
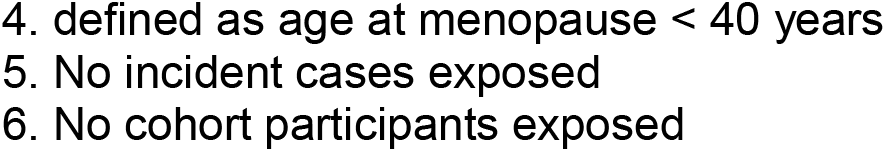
Associations between indicators of endogenous estrogen exposure and incident dementia (adjusted^1^ subhazard ratios with 95% confidence intervals)

| Site | Age at Menarche (per year) | Age at menopause (per year) | Reproductive period (per year) | Parity (per child)<sup>2</sup> | Nulliparity<sup>2</sup> | Age at birth of first child | Index of endogenous estrogen exposure<sup>3</sup> (per SD) | Premature ovarian failure<sup>4</sup> |
| --- | --- | --- | --- | --- | --- | --- | --- | --- |
| Cuba | 1.005<br>(0.916-1.101) | 0.998<br>(0.968-1.030) | 0.998<br>(0.968-1.029) | 0.98<br>(0.89-1.08) | 1.07<br>(0.66-1.73) | 1.008<br>(0.975-1.042) | 1.012<br>(0.925-1.108) | 1.07<br>(0.57-2.00) |
| Dominican Republic | 0.958<br>(0.863-1.063) | 0.996<br>(0.965-1.029) | 1.000<br>(0.968-1.034) | 1.05<br>(1.00-1.10) | 0.66<br>(0.31-1.41) | 0.973<br>(0.934-1.015) | 0.967<br>(0.874-1.069) | 1.53<br>(0.92-2.53) |
| Peru urban | 0.959<br>(0.733-1.255) | 0.986<br>(0.935-1.041) | 0.988<br>(0.932-1.048) | 1.12<br>(0.99-1.26) | Not estimated<sup>5</sup> | 0.963<br>(0.855-1.084) | 0.882<br>(0.744-1.047) | 1.16<br>(0.39-3.42) |
| Peru rural | 1.062<br>(0.772-1.462) | 1.050<br>(0.926-1.189) | 1.040<br>(0.928-1.166) | 1.09<br>(0.92-1.29) | Not estimated<sup>6</sup> | 0.996<br>(0.904-1.098) | 0.953<br>(0.768-1.182) | 1.58<br>(0.44-5.72) |
| Venezuela | 0.924<br>(0.820-1.042) | 0.976<br>(0.940-1.014) | 0.984<br>(0.946-1.024) | 1.04<br>(0.97-1.12) | 0.34<br>(0.05-2.35) | 0.977<br>(0.939-1.015) | 0.975<br>(0.906-1.049) | 1.40<br>(0.63-3.08) |
| Mexico urban | 1.105<br>(0.903-1.353) | 1.007<br>(0.939-1.079) | 0.999<br>(0.935-1.068) | 1.02<br>(0.93-1.11) | 0.61<br>(0.15-2.56) | 1.021<br>(0.943-1.106) | 0.967<br>(0.828-1.130) | 2.47<br>(0.82-7.51) |
| Mexico rural | 1.070<br>(0.902-1.269) | 1.021<br>(0.970-1.075) | 1.010<br>(0.963-1.059) | 1.00<br>(0.91-1.09) | 1.92<br>(0.57-6.47) | 1.075<br>(1.005-1.149) | 1.082<br>(0.931-1.257) | 0.94<br>(0.32-2.75) |
| China urban | 0.967<br>(0.826-1.132) | 0.996<br>(0.932-1.065) | 1.004<br>(0.947-1.064) | 0.95<br>(0.82-1.11) | 2.43<br>(1.02-5.76) | 1.005<br>(0.950-1.064) | 1.063<br>(0.924-1.223) | Not estimated<sup>5</sup> |
| China rural | 0.908<br>(0.731-1.127) | 1.008<br>(0.909-1.118) | 1.030<br>(0.923-1.149) | 0.98<br>(0.84-1.14) | Not estimated<sup>5</sup> | 1.013<br>(0.889-1.155) | 1.114<br>(0.845-1.469) | Not estimated<sup>5</sup> |
| Puerto Rico | 1.001<br>(0.904-1.108) | 1.009<br>(0.983-1.036) | 1.007<br>(0.981-1.033) | 1.03<br>(0.96-1.11) | 1.48<br>(0.79-2.77) | 0.970<br>(0.927-1.015) | 0.959<br>(0.866-1.062) | 0.64<br>(0.34-1.20) |
| Pooled fixed effect | 0.986<br>(0.944-1.030)<br>0.0% | 1.000<br>(0.986-1.013)<br>0.0% | 1.001<br>(0.988-1.015)<br>0.0% | 1.030<br>(1.002-1.059)<br>0.0% | 1.16<br>(0.86-1.56)<br>30.2% | 0.994<br>(0.978-1.011)<br>7.3% | 0.987<br>(0.951-1.025)<br>0.0% | 1.19<br>(0.91-1.55)<br>0.0% |

1. Controlling for age, education and assets
2. Controlling for age, education, assets, and marital status
3. derived from (age at menopause + waist circumference) – (age at menarche + number of live births + age at first birth), all indicators having first been standardised as z-scores
4. defined as age at menopause < 40 years 5. No incident cases exposed 6. No cohort participants exposed

## Discussion

In the largest prospective cohort study to date we have found quite strong evidence that EEE is not importantly associated with subsequent risk of incident dementia. We also failed to replicate a previously reported interaction between APOE genotype and reproductive period. The precision of our null estimates for the hypothesised main effects of proxy indicators of EEE, observed consistently across diverse settings, exclude the possibility of other than trivial effects. The possible exception is premature ovarian failure, a rare exposure, with an upper confidence interval for the ASHR of 1.55, and with suggestive trends towards positive associations in some Latin American sites. While we did observe an association between greater parity and incident dementia, this seems unlikely to be mechanistically explained by cumulative EEE, since the impact of reproductive period on this pathway would be expected to be much greater. Of note, grand multiparity is linked to increased mortality from both diabetes and cardiovascular diseases[31].

In common with some other studies, we did find evidence for an association between reproductive period and baseline cognitive function, somewhat stronger for animal naming than for the CSI-D composite assessment of cognitive function. However, consistent with other studies the effect sizes were very small, and, in our study, were substantially accounted for by plausible confounders.

### Strengths and weaknesses of the study

Strengths of this study are that associations have been assessed longitudinally, in large population-based dementia-free cohorts, encompassing rural and urban catchment area sites in the Caribbean, Latin America, and China. We used meta-analytical techniques to increase the precision of our estimates. Fixed effect meta-analysis is appropriate, given the negligible heterogeneity for all of the associations studied.

We acknowledge some limitations. First there will have been some misclassification of the recalled exposures, which, given the prospective design, is likely to have been non-differential with respect to the outcome. The effect would therefore be towards an attenuation of any genuine association towards the null. Second, we did not gather information on use of hormone replacement therapy (exogenous estrogen). However, availability, awareness and use of such medication can be safely assumed to have been negligible in these countries over the relevant period [16,32-34], particularly in the predominately socio-economically disadvantaged catchment area populations studied. Nevertheless, we cannot exclude the possibility that use of exogenous estrogen could have masked associations with proxy indicators for EEE, particularly if this was used selectively by those experiencing earlier menopause. Third, we did not enquire whether menopause was surgically induced or naturally occurring. Oopherectomy, because of the sudden fall in estrogen levels, might have a particularly marked impact on cognitive functioning[8]. Fourth, it is likely that reproductive history predicts post-reproductive mortality, with a higher mortality risk for those with younger age at first birth, and a U-shaped relationship with parity [35]. To the extent that such associations might selectively remove those who might be at risk for developing dementia from the at risk population, this could bias estimates of association. This possibility is addressed, in part, through our use of competing risk regression to model associations, but this would only account for selective mortality patterns over the follow-up period.

### Contextualisation with other research

The only previous longitudinal study of these associations was in a smaller and younger cohort from Rotterdam (167 incident dementia cases, compared with 692 in our study)[15]. The greater power and precision of our study may not entirely explain the discrepancy in findings; the Dutch study reported a statistically significant increased risk of dementia concentrated in the three-quarters of the cohort with the longest reproductive periods, but only for non-carriers of the APOE e4 genotype. Although the distribution of reproductive periods was similar between the two cohorts, the reproductive history of the Dutch women was very different. Mean parity was 2.2 compared with 4.1 in our study, 11% reported ever using HRT, and 24% reported an artificial menopause from surgery, drugs or radiation therapy.

## Conclusions

We found no evidence to support the theory that natural variation in cumulative exposure to endogenous estrogens across the reproductive period influences the incidence of dementia in late life. Any beneficial effect on cognitive reserve is likely to be very small, and may arise from confounding by shared developmental antecedents. The case for post-menopausal hormone replacement is currently controversial, with conflicting evidence, and some clear risks associated with longer-term use[36]. Nevertheless, the concept of a ‘critical window’ in the immediate post-menopausal period has been widely discussed, during which estrogen replacement therapy may be both less risky, and more beneficial to cognition[8,36,37]. Our study provides only indirect evidence to inform this debate, since our focus was upon pre-menopausal endogenous exposure. However, associations with indicators of endogenous exposure are routinely presented as ‘proof of concept’ for the estrogen hypothesis, and this evidence is significantly weakened by the current study.

## Authors’ contributions

All of the authors (MP, DA, MG, YH, IZJ-V, JJLR, AS, ALS, K-CC, MED, ZL, RM, and AV) worked collectively to conceive the study, and develop the research hypotheses, protocols and methods described in this paper. JJLR (Cuba-Havana), AV (Cuba-Matanzas), DA (Dominican Republic), IZJ-V (Puerto Rico), MG (Peru), AS (Venezuela), ALS (Mexico), and YH (China, assisted by ZL) were principal investigators responsible for the field work in their respective countries. MP conducted the analyses and prepared the first draft of the manuscript. All authors reviewed the draft and provided further contributions and suggestions. All authors had full access to all the data in the study, and read and approved the final report. All authors and the corresponding author had final responsibility for the decision to submit for publication.

## Abbreviations

ASHR: Adjusted Sub-Hazard Ratio APOE-Apolipoprotein E
CERAD: Consortium to Establish a Registry for Alzheimer’s Disease CI-Confidence interval
CSI-D: Community Screening Instrument for Dementia
DSM: Diagnostic and Statistical Manual
EEE: Endogenous estrogen exposure
ICEEE: Index of cumulative endogenous estrogen exposure
POF: Premature ovarian failure

## Availability of supporting data

Data are freely available from the 10/66 Dementia Research Group public data archive for researchers who meet the criteria for access to confidential data. Information on procedures to apply for access to data is available at https://www.alz.co.uk/1066/1066_public_archive_baseline.php, or by contacting Prof Martin Prince at.

## Acknowledgements

### Authors’ information

Martin Prince received salary support from the National Institute for Health Research (NIHR) Mental Health Biomedical Research Centre at South London and Maudsley NHS Foundation Trust and King’s College London.

